# A gene expression analysis and immune infiltration between lesioned and preserved subchondral bone in osteoarthritis

**DOI:** 10.1101/2022.09.12.507705

**Authors:** Gang Zhang, Chengliang Yin, Rilige Wu, Ren Wang, Yong Qin, Songcen Lyu

## Abstract

**Background:** Osteoarthritis (OA) is a degenerative disease which need more research. The purpose of this study was to performed gene expression analysis and immune infiltration between lesioned and preserved subchondral bone, and validation with datasets and experiment of multiple tissues.

**Methods:** The different expressed genes(DEGs) of GSE51588 datasets between lesioned and preserved tibial plateaus of OA patients were conducted. Moreover, functional annotation and protein–protein interaction (PPI) network were applied for exploring the potential therapeutic targets in OA subchondral bones between lesioned and preserved sides. In addition, multiple tissues were used to screen out co-expressed genes and the expression levels of identified candidate DEGs was detected by quantitative real-time polymerase chain reaction (qRT-PCR) in OA. Finally, immune infiltration analysis was conducted.

**Results:** A total of 1010 DEGs were identified, including upregulated 423 genes, 587 downregulated genes. Upregulated genes showed that BP terms were enriched in “skeletal system development”, “sister chromatid cohesion”,”ossification” etc. Pathways were enriched in “Wnt signaling pathway,” “Proteoglycans in cancer,” Downregulated genes showed that BP terms were enriched in “inflammatory response,” “xenobiotic metabolic process”, “positive regulation of inflammatory response”. Pathways were enriched in “Neuroactive ligand–receptor interaction”, “AMPK signaling pathway” etc. JUN, TNF, IL1B,were the hub genes in the PPI network. Col1A1 and LRRC15 were screened out by multiple datasets and validated by experiments. Immune infiltration showed adipocytes and endothelial cells infiltrated less in lesioned samples.

**Conclusion:** Our research might provide valuable information for exploring the pathogenesis mechanism of OA and identifying the potential therapy targets for OA diagnosis.

## Introduction

Osteoarthritis (OA) is a common degenerative and debilitating joint disease. It has become a leading cause of disability and impaired quality of life in the elderly. OA is considered to be an organ disease that affect the whole joint, not only including cartilage, but also including subchondral bone, meniscus, synovium, and ligament[1]. It’s pathology include cartilage loss, synovial hyperplasia, ligament fibrosis, osteophyte formation, subchondral bone remodeling and sclerosis, and increased cytokine production[2]. Subchondral bone plays a crucial role in the pathological process of OA and is an important source of pain. Xu Cao, et al. has demonstrated that osteoclasts derived from subchondral bone can induce sensory innervation and osteoarthritis pain, and alendronate can inhibit osteoclast activity, alleviate aberrant subchondral bone remodeling, reduce innervation and improve pain behavior simultaneously at the early stage of OA[3]. But its mechanism has not been elucidated distinctly.

In the era of “bigdata”, large quantities of data have been accumulated, including bioinformatics[4]. High-throughput sequencing data has developed rapidly and made great contributions in the fields of molecular mechanism, and discovery of drug target. Previous studies have demonstrated gene level changes play important roles in OA diagnosis and development[5, 6]. However, pathological changes are different between OA sides and preserved sides in accordance with cartilage changes, as well as subchondral bone changes[7]. And local therapy and precision medicine of OA need more research.

In this study, we analysis gene expression profiles between lesioned and preserved tibia in OA patients, discovering associated genes, pathways and immune cells, validation with multiple tissues datasets and experiments.

## Results

### Analysis of differential gene expression of all datasets

The DEGs of GSE51588 MT-LT were 1010 genes, including 423 up-regulated genes and 587 down-regulated genes. The most up-regulated gene was STMN2 and POSTN, and the most down-regulated gene was LEP and APOB. The distribution of all DEGs according to the two dimensions of -log10(p value) and logFC is represented by a volcano map in Figure 1A. The DEGs were evaluated by a heatmap, as shown in Figure 1B. DEGs in details with logFC and P value were shown in supplementary 1.

**Figure 1.**
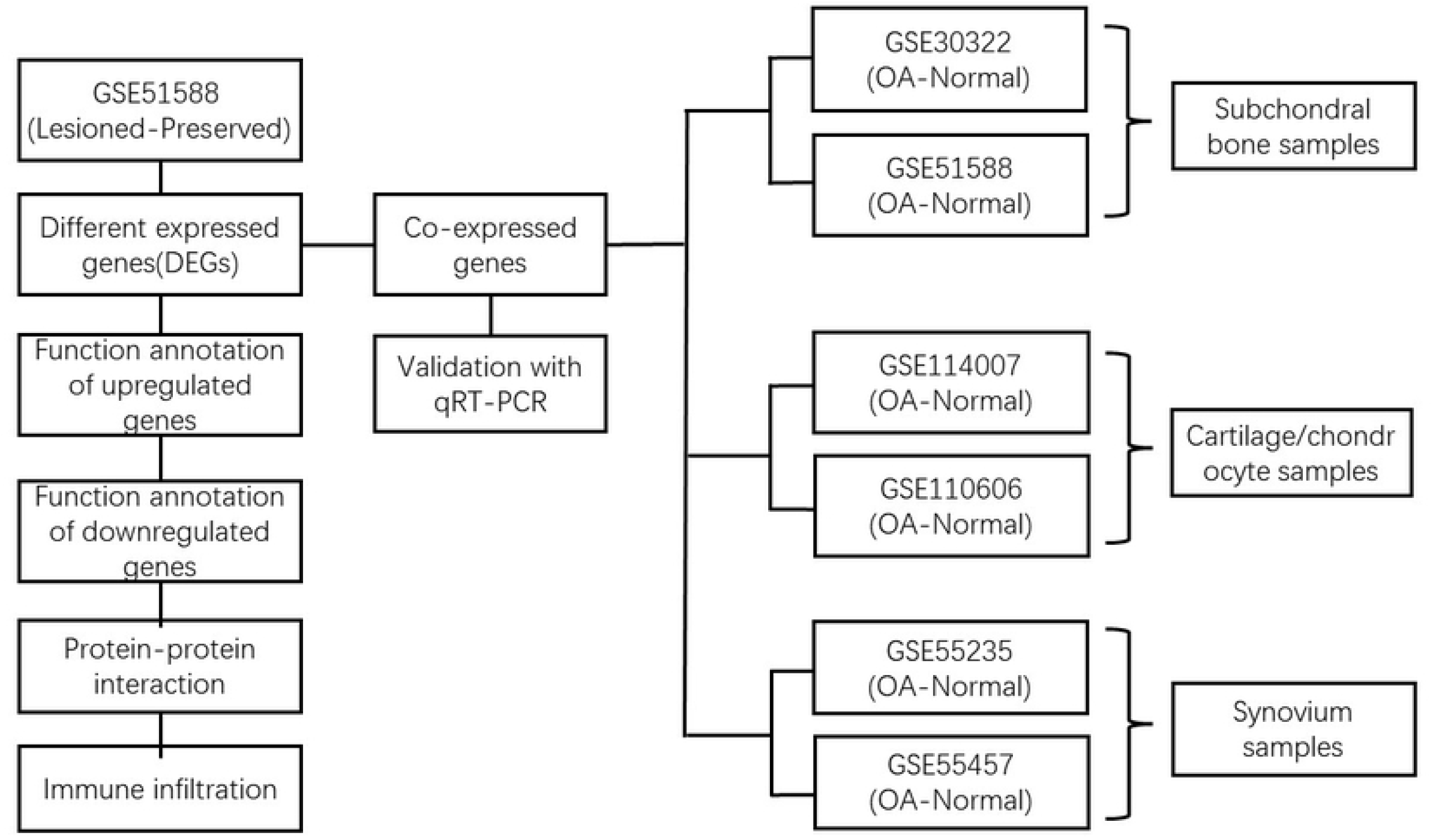
The flow chart depicting the study process for osteoarthritis(OA) gene expression analysis.

### Functional and pathway analysis of DEGs

Functional enrichment analysis of the upregulated DEGs demonstrated that the top five biological processes mainly included skeletal system development, sister chromatid cohesion, ossification, mitotic nuclear division, extracellular matrix organization(Figure 2A). The main cellular components and molecular functions involved in spindle microtubule, proteinaceous extracellular matrix, platelet-derived growth factor binding, metalloendopeptidase activity ect.(Figure 2B and C). KEGG pathways of the upregulated DEGs showed that the top five terms were associated with Wnt signaling pathway, Proteoglycans in cancer, Protein digestion and absorption, PI3K-Akt signaling pathway, Hippo signaling pathway(Figure 2D).

**Figure 2.**
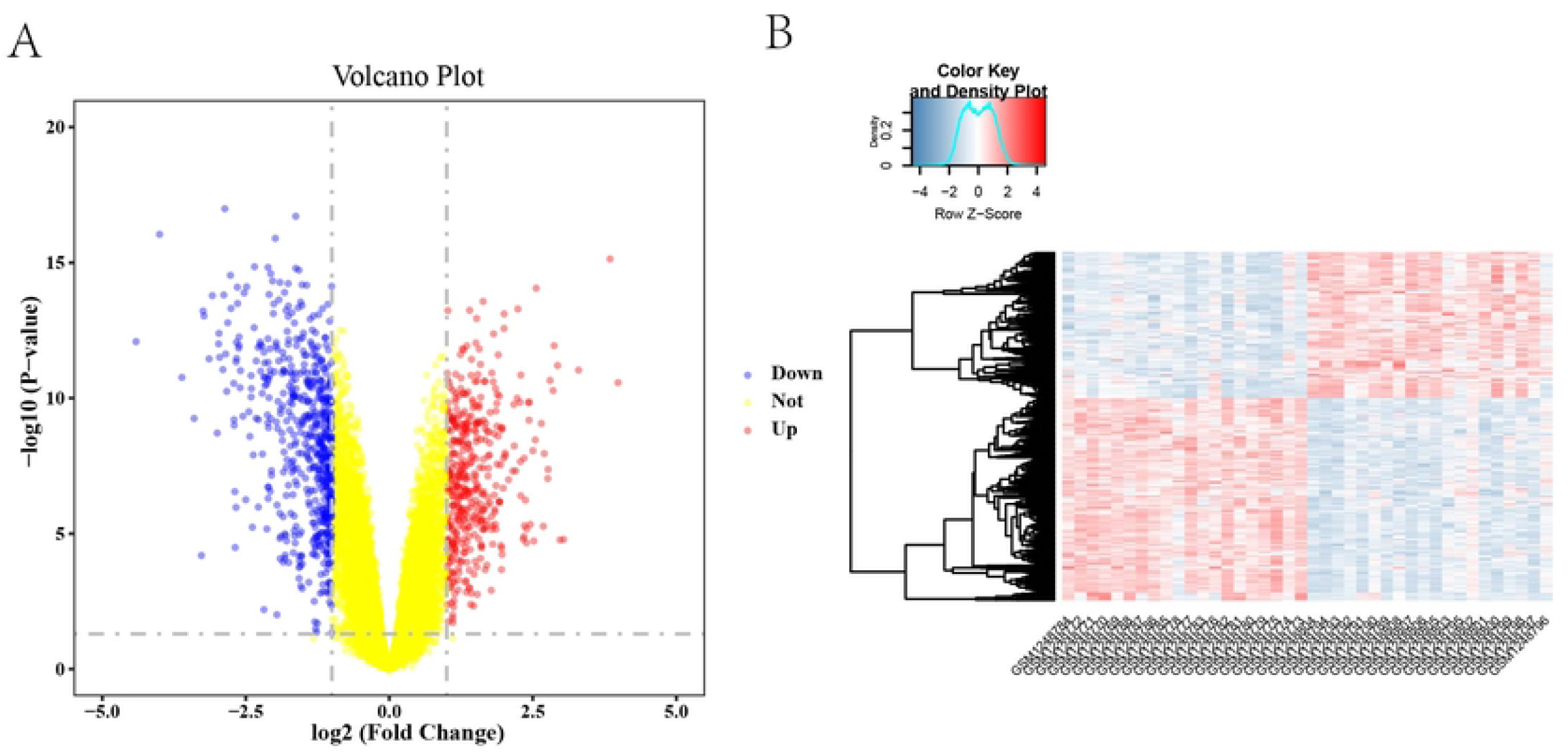
Differentially expressed genes in lesioned and preserved subchondral bone tissues. (A) volcano map of DEGs. The red represents upregulated genes while blue represents downregulated genes. (B) heatmap of DEGs.

The functional enrichment analysis of the downregulated DEGs revealed that the top five BP terms were associated with xenobiotic metabolic process, triglyceride biosynthetic process, positive regulation of inflammatory response, positive regulation of fat cell differentiation, positive regulation of B cell activation(Figure 3A). The main cellular components and molecular functions were associated with receptor complex, plasma membrane, transporter activity, retinol dehydrogenase activity and so on(Figure 3 B and C). KEGG pathways of the down-regulated DEGs showed that the top terms were associated with Tyrosine metabolism, Regulation of lipolysis in adipocytes, PPAR signaling pathway, Neuroactive ligand–receptor interaction, AMPK signaling pathway (Figure 3D).

**Figure 3.**
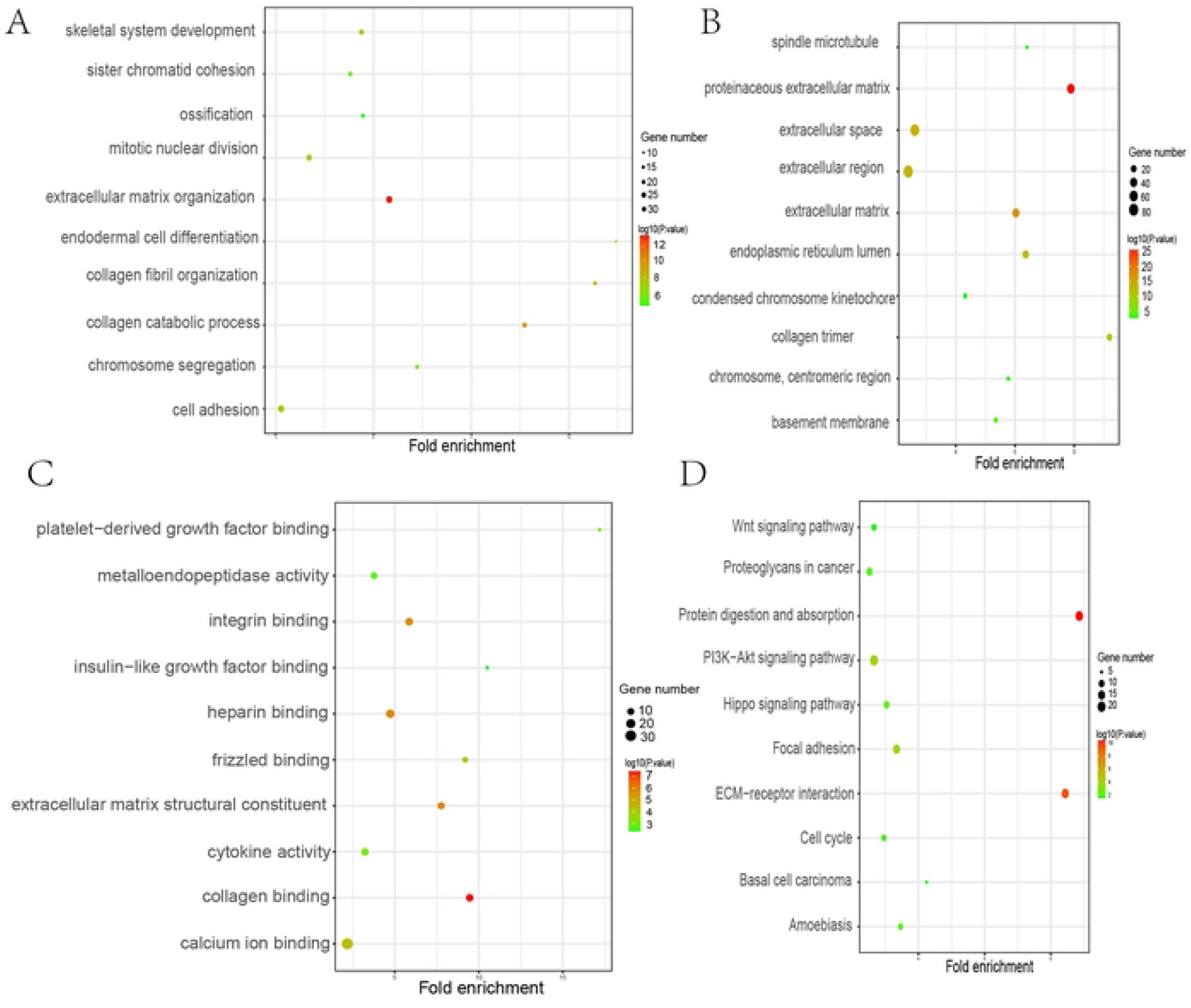
Function annotation of upregulated genes. (A) Biological process analysis; (B) Cellular component analysis; (C) Molecular function analysis. (D) Kyoto Encyclopedia for Genes and Genomes (KEGG) pathway analysis.

### PPI network construction and hub genes identification

To observe the relationships between the DEGs, the PPI network was created using the STRING website. Finally, a total of 1010 DEGs were mapped to 5766 nodes in the network with a combine score>0.4. Network interactions with a score> 0.99, including 116 nodes were visualized with Cytoscape software(Figure 4A). And the top 10 hub genes calculated by MCC were JUN, TNF, IL-1β, LEB, CXCL8, FN1 and so on.(Figure 4B).

**Figure 4.**
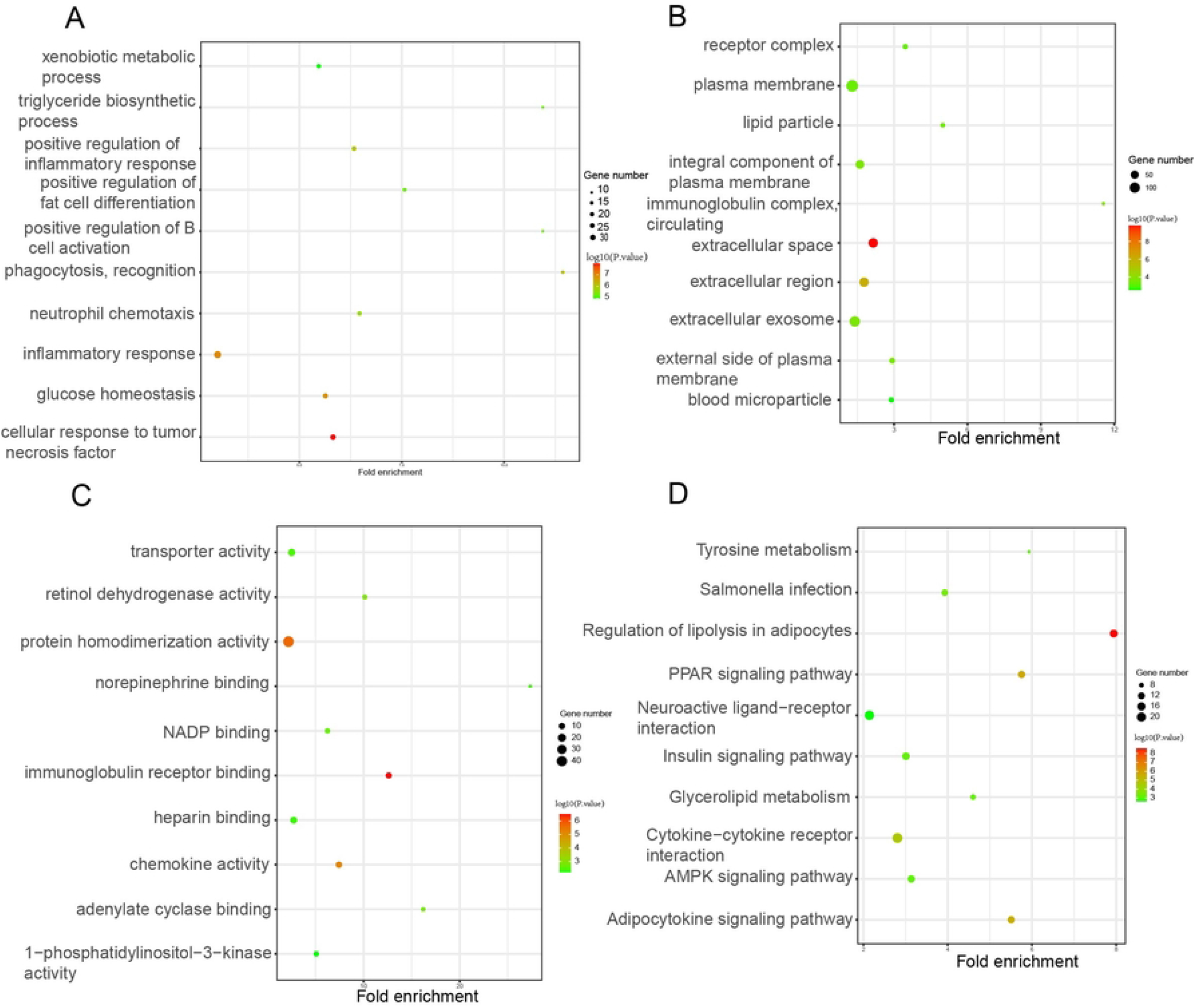
Function annotation of downregulated genes. (A) Biological process analysis; (B) Cellular component analysis; (C) Molecular function analysis. (D) Kyoto Encyclopedia for Genes and Genomes (KEGG) pathway analysis.

### Identification of the co-expressed genes with datasets of multiple tissues and validation with qRT-PCR

Six datasets of multiple tissues including subchondral bone samples, cartilage, and synovium samples were used for validation. Two co-expressed genes were screened out and they were LRRC15 and Col11A1. The results were shown as venn diagram in Figure 5. They were upregulated in all datasets. qRT-PCR was conducted to validate the expression of the co-expressed genes with chondrocyte, FLS, and subchondral bone samples. LRRC15 and Col11A1 were upregulated in all tissues.(Figure 6)

**Figure 5.**
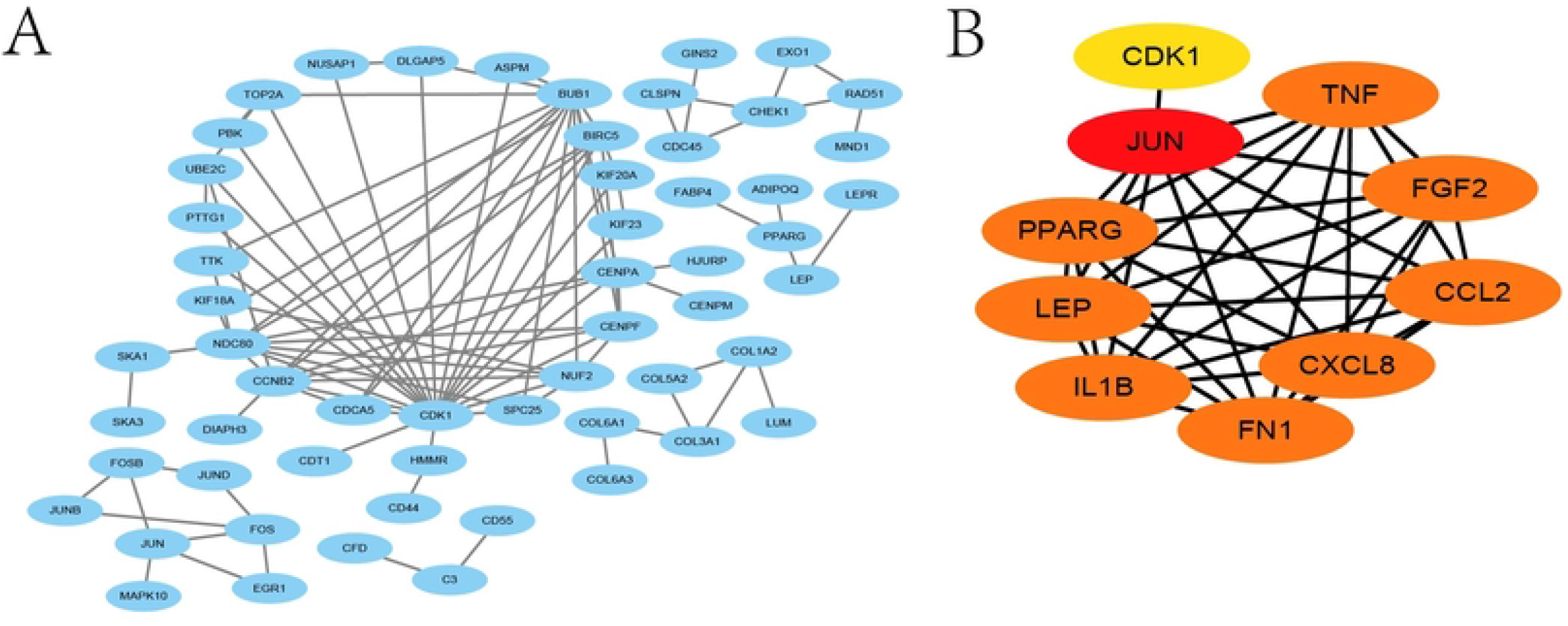
PPI network of DEGs in lesioned and preserved subchondral bone tissues. (A) the combined score was 0.99.(B) The top 10 hub genes showing by MCC method

**Figure 6.**
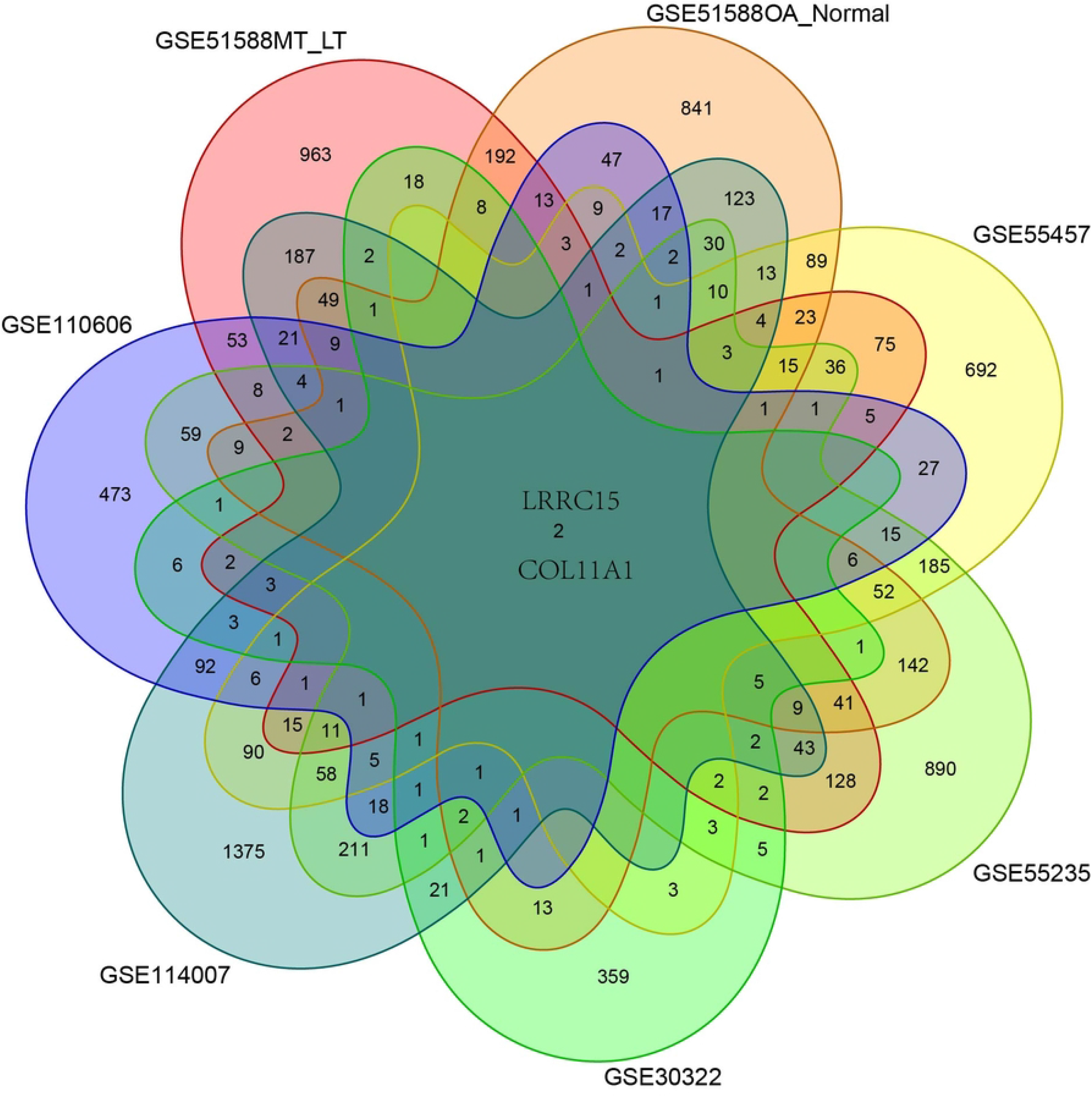
Screen out co-expressed genes with multiple tissues and LRRC15 and Col11A1 are co-expressed genes.

**Figure 7.**
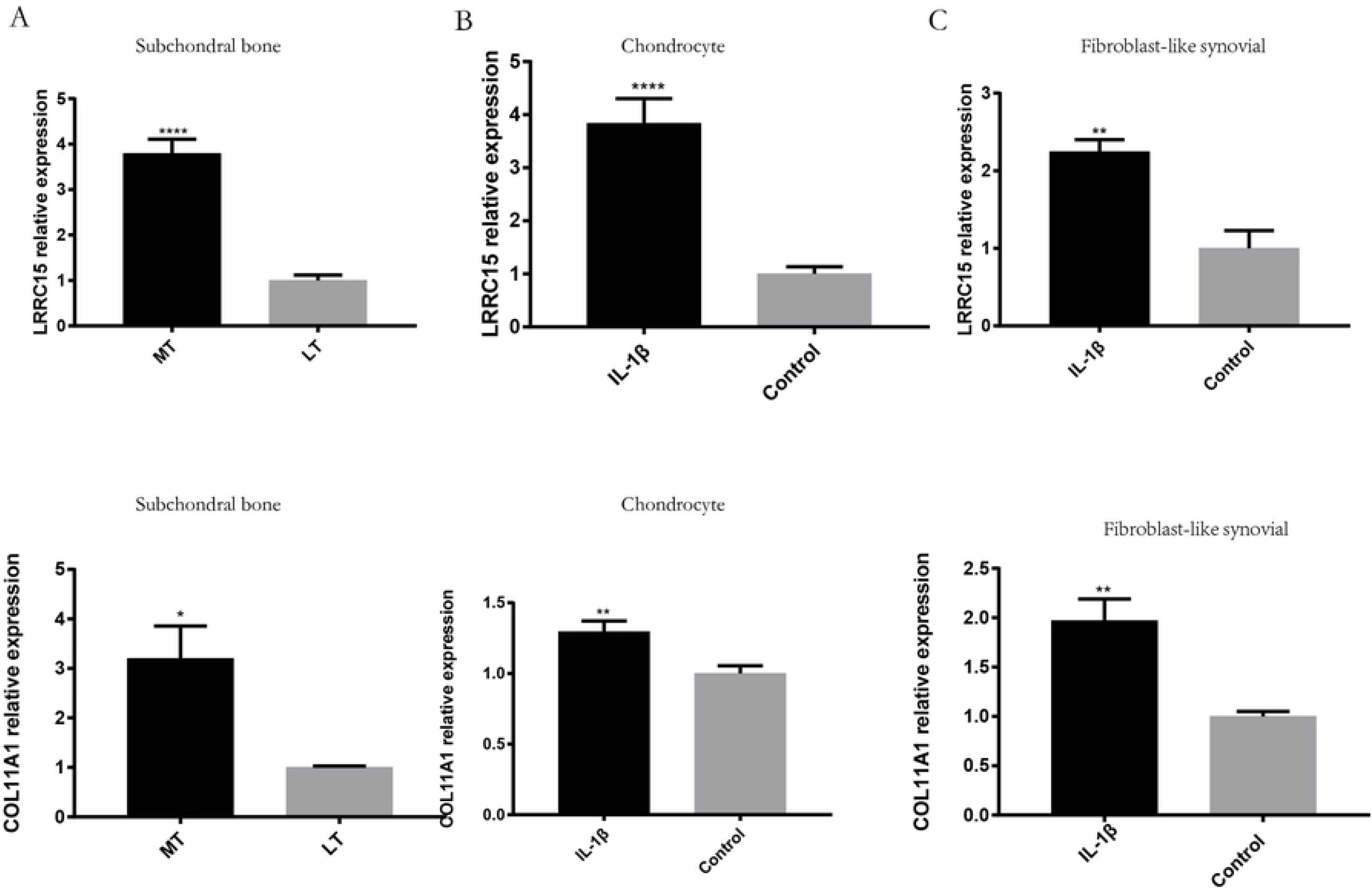
Validation co-expressed genes in multiple tissues with qRT-PCR experiment.

### Analysis of immune infiltrating cells

We performed immune infiltrating cells using xCell algorithm. Correlation heatmap of the 64 types of immune cells revealed that ly Endothelial cells, mv Endothelial cells, and Endothelial cells had a significant positive correlation. CD4 T+ cells and CD4 +Tem also had a positive correlation. (Figure 8A). The heatmap showed abundance score of the immune cells in each sample (Figure 8B).

**Figure 8.**
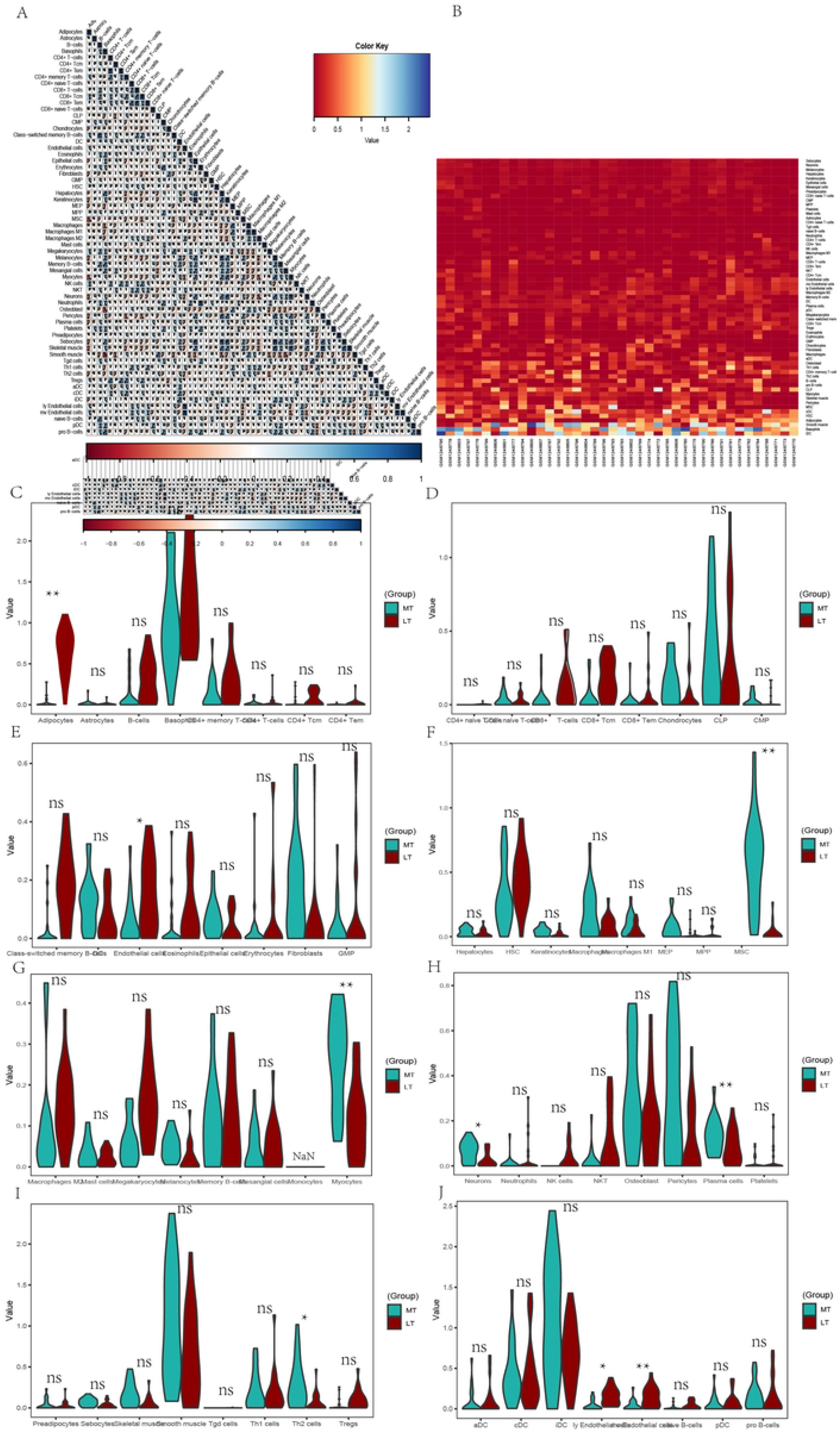
Immune infiltration analysis between lesioned and preserved subchondral bone tissues.(A) The correlation of immune cells. (B) The abundance score of immune cells in subchondral bone samples.(C-J) The difference of immune infiltration between lesioned and preserved subchondral bone tissues.

The violin plot of the immune cell infiltration difference showed that, compared with the preserved control sample, adipocytes and endothelial cells infiltrated less, while msc, myocytes, plasma cells, Th2 cells, ly endothelial cells, mv endothelial cells infiltrated more.(Figure8C-J)

## Discussion and conclusion

In the current study, we identified dysregulated genes associated with OA progression from lesioned and preserved subchondral bone of osteoarthritis. STMN2 and POSTN was most upregulated, and LEP and APOB was most downregulated. The most enriched pathways of upregulated genes was Wnt signaling pathway, while The most enriched pathways of downregulated genes was Tyrosine metabolism. JUN, TNF, IL-1β, LEB was hub genes. LRRC15 and Col11A1 were upregulated and co-expressed in multiple OA tissues. In addition, immune infiltration showed that many immune cells had different infiltrated abundance score.

Previous studies emphasized on articular cartilage degeneration and ignored the role of subchondral bone and synovium. In recent years, more research demonstrated that knee joint is an organ and OA can affect subchondral bone, synovium and other tissues[1, 2, 8]. Subchondral bone links the joint to the diaphyseal bone, provide mechanical support for joint, provide some nutrition and remove metabolic waste products[9]. Subchondral bone remodeling already occurs in early stages of cartilage degeneration. The earliest signs of OA seen on MRI scanning in the subchondral bone are bone marrow lesions (BML; excessive water signals in bone). They are considered to be subchondral bone remodelling due to mechanical overload[10]. The pathological changes of subchondral bone during OA including angiogenesis, de novo bone formation, sensory innervation invasion, bone cysts, sclerosis and osteophytes formation[3, 11, 12]. More evidence suggests that abnormal bone remodeling may contribute to the development of OA, and can be a target for OA therapy [13-15].

The study identified the most up-regulated gene was STMN2 and POSTN, and the most down-regulated gene was LEP and APODB. STMN2(stathmin2) is associated with sensory neurons growth and contribute to regenerating axons after nerve injury[16].Maybe it is implicated in the pain of OA. POSTN (periostin) is a 90-kDa member of the fasciclin family. POSTN can induce the expressions of proinflammatory factors, such as MMP-9, MMP-10, and MMP-13 production, leading to degradation of the extracellular matrix[17]. LEP(*Leptin*)is a peptide hormone containing 167 amino acids [18]. LEP promotes the differentiation of osteoblast under normal conditions[19]. LEP expression was increased in OA subchondral osteoblasts and in part elevated the expression levels of alkaline phosphatase, osteocalcin release, collagen1 and TGF-β1. But in OA conditions, osteoblast, osteoclast and subchondral bone remodeling have disturbed [20]. And further research needs to be performed.

For upregulated DEGs, the most enriched pathway was enriched in Wnt signaling pathway, PI3K-Akt signaling pathway and so on. Wnt signaling pathway can directly affect subchondral bone cartilage and synovial tissue, which has been proven to play important roles in pathology[21, 22]. Phosphorylation of AKT in subchondral bone can promote osteogenic differentiation and osteoblastic proliferation, and resulting in aberrant bone formation. And targeting these pathways maybe alleviate the development of OA. By contrast, inhibition of PI3K/AKT reduces subchondral bone sclerosis through decreasing osteogenesis[23]. Also suppressing PI3K-Akt signaling pathway can enhance cell autophagy, reduce chondrocytes inflammation[24]. And Pharmaceutical intervention of the pathway provide a promising approach for OA treatment[25].

For downregulated DEGs, the most enriched pathways include Tyrosine metabolism, AMPK signaling pathway and so on. AMP activated protein kinase (AMPK) mainly regulate energy balance and metabolism. Dysregulation of AMPK is associated with multiple age-related diseases including atherosclerosis, cardiovascular disease diabetes, cancer, neurodegenerative diseases and OA[26, 27]. Upregulated of Phosphorylation and total AMPK expression in articular cartilage can limit OA development and progression in OA animal models[28, 29].

10 hub genes were identified in the PPI network, and the proteins encoded by these genes are the key nodes in the PPI network, including JUN, TNF, IL-1β, LEB, CXCL8, FN1 and so on. They are associated with OA progression, especially JUN, IL-1β and TNF[30-32]. JUN is a major component of the transcription factor activator protein-1 (AP-1)family[33]. As a transcription factor, JUN can mediate catabolic transcription and cell apoptosis/death, also play an important role in the TNF signaling pathway and IL-17 signaling pathway [34].

With other datasets of multiple tissues and experiments, LRRC15 and COL11A1 were identified as co-expressed genes. These two novel genes are proposed to play important roles in the pathogenesis of OA. LRRC15, type I membrane protein, has 581 amino acids, with no obvious intracellular signaling domains, is upregulated by the pro-inflammatory cytokine TGFβ in cancer-associated fibroblasts [35]. It is highly expressed both on stromal fibroblasts as well as tumor cells, such as melanoma sarcomas, and glioblastoma[36]. ABBV-085 is an antibody drug, which can directedly against LRRC15. At present, it has been widely studied for antitumor research. [37]. Maybe, it can be applied in OA treatment. Otherwise, LRRC15 was significantly downregulated in OVX mice and upregulated upon osteogenic induction in a p65-dependent manner[38]. Collagen XI which encodes the α1 chain of type XI collagen, is essential for collagen fibril formation in articular cartilage[39]. Primarily type II collagen with the alpha 1collagen XI [α1(XI)] chain structured into collagen fibrils, form a network in the cartilage extracellular matrix (ECM) that contributes to retention of proteoglycans and tensile strength in cartilage tissue[40].And many studies have shown that *COL11A1* may have an increase the genetic susceptibility to develop OA[41-43].

The inflammation and angiogenesis can alter the process of subchondral bone modelling[8]. By analyzing the immune infiltration, it was found that ly Endothelial cells, mv Endothelial cells, and Endothelial cells had a significant positive correlation. CD4 T+ cells and CD4 +Tem also had a positive correlation. Adipocytes and endothelial cells infiltrated less in lesioned samples, while msc, myocytes, plasma cells, Th2 cells, ly endothelial cells, mv endothelial cells infiltrated more in lesioned samples.

In conclusion this study suggests that Subchondral bone has important roles in the progression of OA. Understanding the mechanisms of subchondral bone involved in OA development and progression will greatly contribute to the diagnosis, treatment, and prognosis of OA patients.

## Materials and methods

### Data processing and identification of DEGs

The NCBI Gene Expression Omnibus(GEO) is a public data repository that stores gene expression profiles, raw series and platform records (https://www.ncbi.nlm.nih.gov/geo/). Subchondral bone dataset GSE51588 included 10 normal datasets, and 40 OA samples. The platform was <u>GPL13497.</u> The 40 OA samples included medial tibia samples (significant degeneration, lesioned) and lateral tibia samples (minimal degeneration, preserved). And the OA samples were used for further training analysis. The data matrix series were downloaded and the limma package were used in R software to identify DEGs[44]. The *p* < 0.05 and |logFC2|>1was considered as the cutoff criterion.

### Functional annotation of DEGs

The Database for Annotation, Visualization, and Integrated Discovery (DAVID; https://david.ncifcrf.gov/) provides a comprehensive set of functional annotation tools and help investigators to understand biological meaning behind large list of genes. Gene Ontology (GO) annotation consists of cell components (CCs), molecular functions (MFs), and biological processes (BPs) of genes. The KEGG pathway enrichment analysis is used to extract pathway information from molecular interaction networks. In the present study, the DAVID online tool was used to conduct GO and KEGG pathway enrichment analysis of up-regulated and down-regulated genes respectively. R language was used for data visualization, and *P* < 0.05 was considered to indicate a statistically significant GO and KEGG terms.

### Analysis of PPI network and hub genes

Protein-protein interaction(PPI) networks construction is critical to understanding cell biology and interpreting genomic data. Search Tool for the Retrieval of Interacting Genes/Proteins (STRING) database (http://www.string-db.org/) is an online biological database and website designed to construct the PPI networks in molecular biology. For a more in-depth understanding of the DEGs, PPI network was conducted using STRING database. The DEGs (species: Homo species) were mapped to STRING database and (PPI score >0.4), and the results of combine score>0.99 were shown using Cytoscape software 3.7.1(http://www.cytoscape.org/) to visualize the PPI network, and the top 10 hub genes were calculated by MCC method.

### Screen out co-expressed DEGs using multiple datasets

Six GSE datasets related to joint tissues of OA patients were used for validation to screen co-expressed genes in multi-tissues. The DEGs of associated datasets including GSE51588 OA-Normal for subchondral bone samples,GSE30322 for subchondral bones of SD rats OA models, GSE 110606 and GSE114007 for chondrocyte & cartilage samples, GSE55235 and GSE55457 for synovial tissues were used in this study to screen out co-expressed genes which were in accordance with the expression profile of OA subchondral bone samples. Differentially expressed genes(|logFC2|>1, p < 0.05)were taken forward for further validation. And the results were shown in Venn diagram.

### Quantitative reverse-transcription polymerase chain

Samples from total knee placement were collected. Ethical approval and consent for use of resected tissue were obtained. Subchondral bone samples were obtained and stored in liquid nitrogen until detection. Cartilage and synovium samples were cut into small pieces about 1 mm^3^ and put into collagenase 2 and 1 respectively for about 4 hours. Then α-MEM containing 10% fetal bovine serum and 1% penicillinstreptomycin were added. Three days later, we discarded debris, residual contamination macrophages was avoided three passages later and monocultures of Synovial fibroblasts(SF) were obtained and used for experiments until passage 8[45]. And passage 2 of chondrocyte was used for experiments. SF and chondrocyte were cultured in 6-well plates dishes and 10ng/ml IL-1β were added into SF and chondrocyte for 48h for further research.

Total RNA from SF, chondrocyte and subchondral bone was obtained using reagent(ES science technique). RNA was reverse transcribed using ES science cDNA Reverse Transcription Kit and quantitative real-time PCR analysis was performed with ES science SYBR Green kit. The primers were list in Table 1. For relative quantification, we normalized the target gene expression to the housekeeping gene (GAPDH). Results were presented as the relative expression with respect to the untreated condition using the formula 2-ΔΔCt method. Data was analyzed with GraphPad Prism7.0. Data and differences were assessed using a Student’s two-tailed t-test with P < 0.05 considered significant.

**Table 1.**
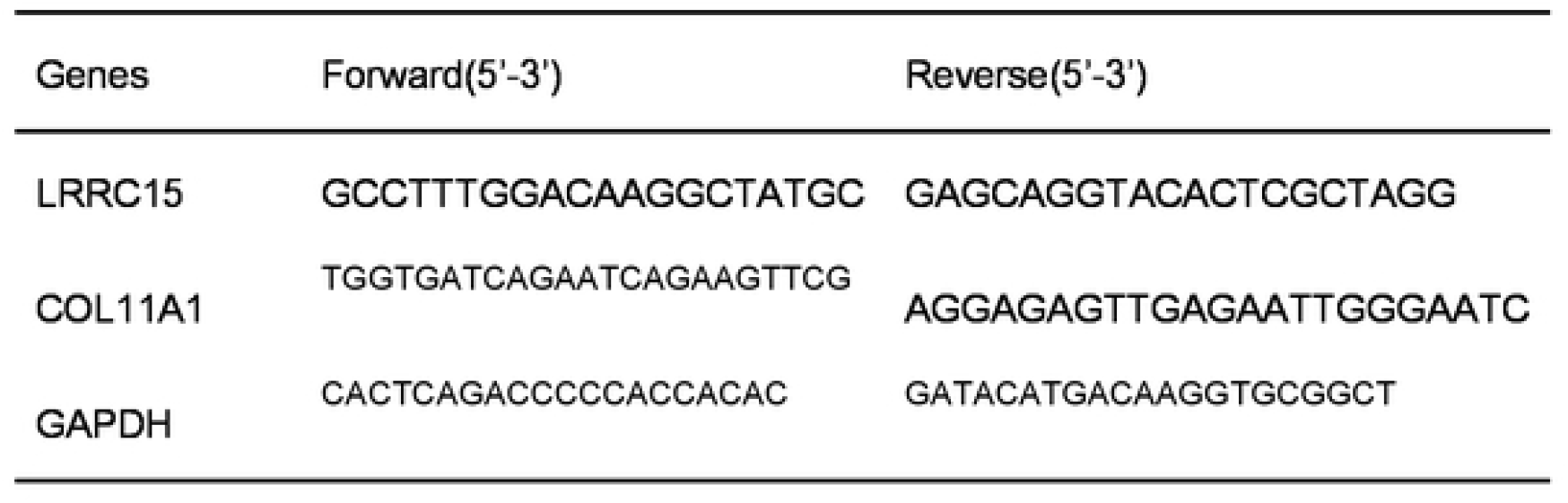
Primers of genes

**Table 2.** The clinical information of patients in datasets.

| Datasets | samples | NO. | GPL<br>platform | SEX(MALE/FEMALE) | Age,<br>years<br>(mean<br>± SD) |
| --- | --- | --- | --- | --- | --- |
| GSE5158 | Subchondral<br>bone |  |  |  |  |
| GSE30322 | Subchondral<br>bone |  |  |  |  |
| GSE114007 | cartilage |  |  |  |  |
| GSE110606 | chondrocyte |  |  |  |  |
| GSE55457 | synovium |  |  |  |  |
| GSE55235 | synovium |  |  |  |  |

| GEO ID | Author | Platform | Samples | Year | Type | Omics |
| --- | --- | --- | --- | --- | --- | --- |
| GSE82107 | Marieke de Vries | GPL 570 | OA: Normal = 10:7 | 2016 | Synovial membrane | mRNA |
| GSE55457 | Woetzel D | GPL 96 | OA: Normal = 10:10 | 2014 | Synovial membrane | mRNA |
| GSE55235 | Woetzel D | GPL 96 | OA: Normal = 10:10 | 2014 | Synovial membrane | mRNA |
| GSE12021 | Huber R | GPL 96 | OA: Normal = 10:9 | 2008 | Synovial membrane | mRNA |
| GSE63359 | Attur M | GPL 96 | OA: Normal = 44:26 | 2017 | Blood | mRNA |
| GSE48556 | Ramos YF | GPL 6,947 | OA: Normal = 106:33 | 2013 | Blood | mRNA |

**Table 2.** The clinical information of patients in datasets.
|  | OA | Normal |
| --- | --- | --- |
| **GSE82107** | 10 | 7 |
| Age, years (mean ± SD) | NA | NA |
| Sex |  |  |
| Male | NA | NA |
| Female | NA | NA |
| **GSE55457** | 10 | 10 |
| Age, years (mean ± SD) | 72.4 ± 5.9 | 51.0 ± 19.7 |
| Sex |  |  |
| Male | 2 | 8 |
| Female | 8 | 2 |
| **GSE55235** | 10 | 10 |
| Sex |  |  |
| Male | 2 | 7 |
| Female | 8 | 3 |
| **GSE12021** | 10 | 9 |
| Age, years (mean ± SD) | 71.9 ± 2.0 | 49.9 ± 6.7 |
| Sex |  |  |
| Male | 2 | 7 |
| Female | 8 | 2 |
| Disease duration, years (mean ± SD) | 6.2 ± 2.7 | 0.4 ± 0.3 |
| CRP(mg/L) | 7.6 ± 2.9 | NA |
| **GSE63359** | 46 | 26 |
| Age, years (mean ± SD) | 65.6 ± 10.7 | 54.6 ± 9.5 |
| Sex |  |  |
| Male | 14 | 7 |
| Female | 32 | 19 |
| BMI (mean ± SD) | 27.4 ± 4.1 | 24.9 ± 3.9 |
| **GSE48556** | 106 | 33 |
| Age, years (mean ± SD) | 58.2 ± 7.6 | 60.9 ± 7.7 |
| Sex |  |  |
| Male | 1 | 9 |
| Female | 105 | 24 |
| **GSE51588** | 40 | 10 |
| Age, years (mean ± SD) | 69.7 ± 8.8 | 38.4 ± 12.7 |
| Sex |  |  |
| Male | 18 | 4 |
| Female | 22 | 6 |

### Computational analysis of immune infiltrating cells

xCell is a novel and robust method based on ssGSEA (single sample Gene Set Enrichment Analysis) that estimates the abundance scores of 64 cell types in the microenvironment. There are two approaches to run xCell, one is the online website tool (https://xCell.ucsf.edu/), the other is xCell package in R software. We estimated the abundance scores of 64 cell types in lesioned and preserved subchondral bone samples using the online website tool. *P* < 0.05 was considered as significant. Also, the immune cell scores were also calculated in each sample. Moreover, the relationship between immune cell were calculated using Pearson coefficient. “ggplot2” package was used to draw violin diagrams to visualize the differences in immune cell infiltration.

## Supplementary Information

The online version contains supplementary material available at

## Authors’ contributions

Yong Qin and Songcen LV designed the research; Gang Zhang collected patient samples and wrote the manuscript; Chengliang Yin and Rilige Wu conducted data mining and analysis. Ren Wang conducted performed experiments validation. All authors read and approved the final manuscript.

## Declarations

### Ethics approval and consent to participate

This study was approved by the Ethics Committee of the second Affiliated Hospital of Harbin Medical University (Approval number: ky2020-078) and informed consent was taken from all the patients.

### Consent for publication

All authors consent to publication.

